# Subnuclear localisation is associated with gene activation not repression or parental origin at the imprinted *Dlk1-Dio3* locus

**DOI:** 10.1101/2021.03.17.435843

**Authors:** Rahia Mashoodh, Lisa C. Hülsmann, Frances L. Dearden, Nozomi Takahashi, Anne C. Ferguson-Smith

**Author notes:** Corresponding author, (ACFS). These authors contributed equally to the work (Rahia Mashoodh, Lisa C. Hülsmann).

## Abstract

At interphase, de-condensed chromosomes have a non-random three-dimensional architecture within the nucleus, however, little is known about the extent to which nuclear organisation might influence expression or *vice versa*. Here, using imprinting as a model, we use 3D RNA- and DNA-fluorescence-in-situ-hybridisation in normal and mutant mouse embryonic stem cells to assess the relationship between imprinting control, gene expression and allelic distance from the nuclear periphery. We compared the two parentally inherited imprinted domains at the *Dlk1-Dio3* domain and find a small but reproducible trend for the maternally inherited domain to be further away from the periphery if the maternally expressed gene *Gtl2/Meg3* is active compared to when it is silenced. Using Zfp57KO ES cells, which harbour a paternal to maternal epigenotype switch, we observe active alleles significantly further away from the nuclear periphery with the distance from the periphery being proportional to the number of alleles active within the cell. This distribution of alleles suggests an activating effect of the nuclear interior rather than a repressive association with the nuclear periphery. Although we see a trend for the paternally inherited copy of the locus to be closer to the nuclear periphery, this appears to be linked to stochastic gene expression differences rather than parental origin. Our results suggest that transcriptional activity, rather than transcriptional repression or parental origin, defines sub-nuclear localisation at an endogenous imprinted domain.

**Author summary:** Genomic imprinting is an epigenetically regulated process that results in the preferential expression of a subset of developmentally regulated genes from maternally or paternally inherited chromosomes. We have used imprinted genes as a model system to investigate the relationship between the localisation of genes within the cell nucleus and their active expression while at the same time distinguishing gene repression by genomic imprinting, and gene repression by other mechanisms that act on the active allele. We find that there is a significant correlation between transcription and distance to the edge of the nucleus for the *Gtl2/Meg3* gene in the imprinted *Dlk1*-*Dio3* region. However, this correlation has a very small effect size and the nuclear envelope, which is commonly thought to act as a repressive environment for gene expression, does not appear to play a major role. We show that position effects, which have been shown for artificially lamina-targeted genes, also exist for endogenous loci and consider the possible biological relevance of the observed small effect.

## Introduction

The spatial organization of chromosomes in the interphase nucleus is non-random and involves 3D interactions on chromatin as well as interactions with various nuclear domains, the nuclear envelope being the best characterized (1). While the bulk of chromosomes is arranged as relatively compact and spatially defined chromosome territories (2), open chromatin between these contains transcription factories where active genes from different genomic loci co-localize at times of transcription (3). Recently, there has been much interest in understanding ways in which the 3D localisation of chromatin can influence or be influenced by gene expression (4,5). While chromatin organization into chromosome territories, topologically associated domains, and loops have complex effects on transcriptional regulation, interactions with the nuclear envelope are generally seen as transcriptionally repressive (1). Lamina associated domains (LADs) have been characterized as gene poor, transcriptionally inactive regions (6), even though these repressed regions have been shown to be dynamic in dividing cells and are therefore not always lamina-associated in every cell within a population (7). A notable exception to the repressive environment at the nuclear periphery is represented by nuclear pore complexes, where chromatin-facing nucleoporins have been implicated in transcriptional activation rather than repression in yeast (8). However, in Drosophila, nucleoporins appear to have this activating effect mainly in the nuclear interior (9–11). In line with the general notion that the nuclear periphery acts as a repressive environment, some though not all individual genomic loci have been shown to become transcriptionally repressed after artificial targeting to the nuclear lamina (12–16). These findings suggest a role for subnuclear position in gene regulation, however, the extent and importance of non-random localisation is poorly understood.

The *Dlk1-Dio3* imprinted domain on mouse chromosome 12 is well characterized and is a valuable model for comparative analysis of gene regulatory mechanisms. Imprinted genomic regions contain genes that are preferentially expressed from either the maternally or paternally inherited chromosome, but that also are subject to the same regulated and stochastic transcriptional mechanisms that govern the expression of other genes. Imprinted genes are therefore regulated by both germline-derived parental-origin-specific epigenetic mechanisms and the transcriptional milieu of a particular cell type (17,18). *Dlk1-Dio3* imprinting (Fig 1A) is regulated by an intergenic differentially methylated region (IGDMR) and contains a number of paternally expressed protein-coding genes as well as maternally expressed non-coding RNAs. Recent evidence has explored the relationship between the parental origin of this imprinted domain and its subnuclear locations. Kota and colleagues identified a differential localisation of the *Gtl2/Meg3* gene in a mouse embryonal stem (ES) cell line with the non-expressed paternal copy being closer to the nuclear periphery than the expressed maternal allele (19). Furthermore, a LINE1 (L1) repeat cluster located between *Begain* and *Dlk1* within the cluster (Fig.1A) was described to represent a facultative LAD in mouse ES cell-derived neural stem cells (20,21). Although this suggests that the L1 repeat might have an inhibiting effect on local gene expression via lamina tethering, deletion of the repeat cluster itself did not lead to increased expression of neighbouring genes (20). Here, we use extensive 3D RNA-DNA fluorescence-in-situ-hybridisation (FISH) in multiple normal and mutant mouse ES cell lines and show that gene expression rather than the parental origin of alleles is associated with differences in intranuclear localisation, and suggest that juxtaposition to the nuclear envelope appears to play a minor role, if any, in gene regulation at this endogenous locus.

**Figure 1:**
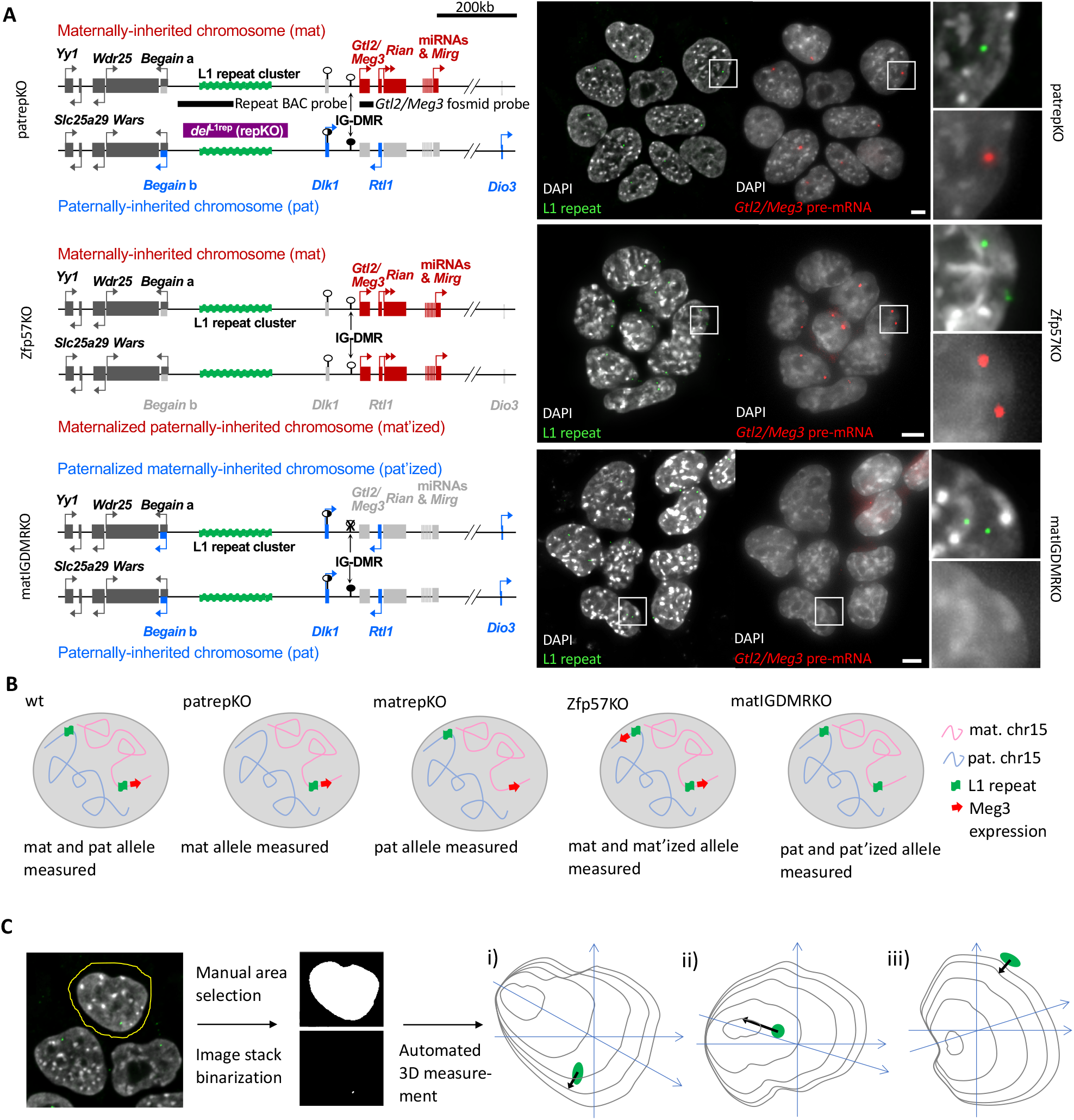
Experimental design.

(**A**) A 260kb deletion encompassing the LINE1 cluster in the *Dlk1-Dio3* imprinted region was used to mark the parental origin of alleles. ES cells derived from mutant mice were used that mimic a uniparental origin of the *Dlk1-Dio3* region. ES cells with a homozygous deletion of the chromatin modifier Zfp57 show maternal expression patterns at the paternally inherited chromosome while ES cells with a maternally inherited deletion of the IGDMR show paternal expression patterns from the maternally inherited chromosome. The FISH images of ES cells, where the maternal L1 cluster was used for 3D measurements and *Gtl2/Meg3* expression was assessed in parallel. As predicted, many Zfp57KO ES cells showed biallelic expression of maternally expressed *Gtl2/Meg3*, while matIGDMRKO ESC showed no *Gtl2/Meg3* expression. Chromosome maps are derived from Soares et al. 2018 (20). The FISH images represent maximum projections of 20-30 central z planes of acquired image stacks. Blow-ups of framed regions are shown on the right. Scale bars represent 50 μm. (**B**) Schematic of nuclei with the expected combined DNA and RNA FISH results for all genotypes. (**C**) Automated, unbiased, distance measurements using the Fiji imaging package. Each nucleus is manually selected from the FISH image stack and both channels are binarized and added to Fiji’s 3D manager to automatically measure the shortest distance from the centre of the FISH signal to the chromatin border in 3D. The shortest distance might be oriented more in the xy axis (i) or more in the z axis (ii) of the image stack. Occasionally, peripheral FISH signals were just outside of the chromatin representation (iii) and therefore measured as negative distances (See Methods).

## Results

### Differential intranuclear distribution of parental alleles of the *Dlk1-Dio3* region

In order to analyse the intranuclear localisation of alleles of the *Dlk1-Dio3* region, while knowing their parental origin, we generated ES cells derived from *del*^L1rep^ maternal and paternal heterozygous mice (20). We used only male ES cell lines for this study since it has been previously shown that imprinting status is more stable in male ES cells compared to female ES cells (22). These cells carry a deletion between the genes *Begain* and *Dlk1*, which includes a 170kb L1 repeat array (Fig. 1A and Supplementary Fig. S1) and have a normal epigenotype. Using a DNA FISH probe specific to the L1 repeat array, we used the deletion to distinguish the two parental chromosomes in 3D DNA FISH experiments, independent of gene expression. Importantly, we only quantified the behaviour of WT chromosomes not carrying the deletion. For each cell we measured the 3D distance of the DNA FISH probe to the nuclear periphery defined by the edge of DAPI staining of chromatin in an automated way (Fig. 1C). This measurement was used as a read-out for the intranuclear position of the *Dlk-Dio3* imprinted region, which lies within a single TAD in ES cells (Supplementary Fig. S1) (23,24). In parallel, we used nascent RNA FISH to obtain the *Gtl2/Meg3* expression state (expressed or non-expressed) for each allele.

We took DNA FISH probe distance measurements in three biological replicates each of heterozygous ES cells derived from maternal inheritance of *del*^L1rep^ (matrepKO) and paternal inheritance of *del*^L1rep^ (patrepKO) and WT ES cells, this way comparing paternally inherited alleles, maternally inherited alleles and the mixture of both as a control. Using a multiple least-squares linear regression model, we find that paternal alleles are closer to the nuclear periphery compared to maternal alleles after controlling for nuclear volume (t=2.946, p=0.00334, estimate (slope) = 0.22; Fig. 2A). Given that WT cells contained both alleles, we decided to characterise the relationship between the two alleles to understand their natural distribution within these cells. In WT cells, one allele was always closer to the nuclear border than the other, and there was a significant difference between the two alleles (t=22.80, p<0.001, estimate (slope)=1.08; Fig. 2A). While the parental origin of the WT alleles was unknown, we assume that this difference might, at least in part, reflect the tendency for the paternal allele to be biased towards the periphery. However, in the KO cells, the average difference in distance between the maternal (*M*=1.81, *SD*=1.00) and the paternal allele (*M*=1.58, *SD*=0.954) was unexpectedly small (230nm on average). Previous work had found differences of more than a micrometer, a difference that is more similar to that between the WT near and far alleles (Fig. 2A) (19). This small difference in KO was not due to variability in replicates as replicates were not significantly different from one another (Supplementary Fig. S2), but suggested that the parental origin effect we observed might be of limited functional significance. We, therefore, developed a genetic approach to compare the two parental chromosomes in a more functional context.

**Figure 2:**
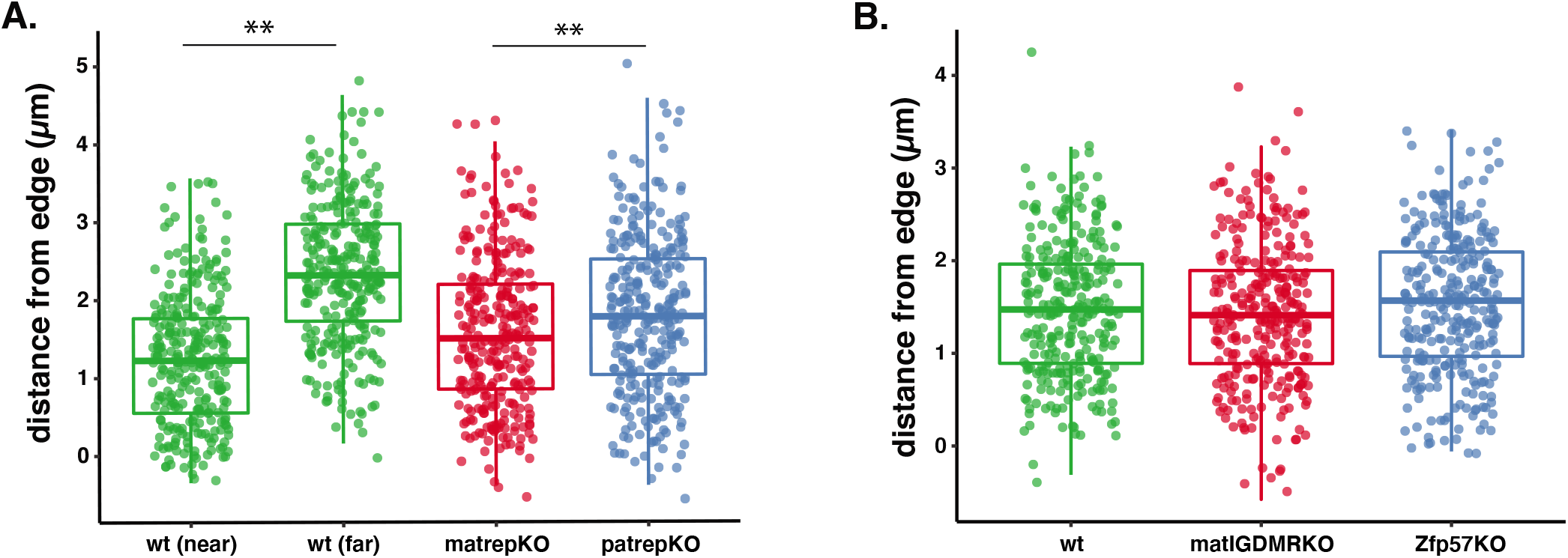
Distribution of alleles by parental origin.

(**A**) Paternal alleles (matrepKO) of the *Dlk1-Dio3* are significantly closer to the nuclear periphery than maternal alleles (patrepKO), as has been reported previously. This difference is less pronounced than the natural difference between the two alleles within a WT cell (\*\**p*<0.01). (**B**) There were no differences in the distance from the periphery between paternal and paternalized alleles from matIGDMRKO ES cells or maternal and maternalized alleles from Zfp57KO ES cells from WT cells.

### Epigenotype switching shifts localisation

To address the relationship between parent of origin and function more overtly we utilized genetic models exhibiting an epigenotype switch that reversed the imprints on either the maternal or the paternal chromosome. Zfp57KO ES cells carry a null mutation in the chromatin modifier ZFP57, which is important for imprint maintenance and, in case of the *Dlk1-Dio3* region, results in a paternal to maternal epigenotype switch causing a maternal expression pattern from the paternally inherited chromosome (Fig. 1A and B and Supplementary Fig. S3) (25,26). Conversely, matIGDMRKO ES cells are derived from blastocysts that have a maternally inherited deletion of the IGDMR, resulting in a maternal to paternal epigenotype switch and expression of the paternally expressed imprinted genes from the maternally inherited chromosome (27). Matched WT control ES cells (WT2) were generated for this experiment. Using three biological replicates for each mutant and control line, we found that there was no significant difference in the subnuclear localisation of mutant chromosomes compared to WT between cell types (t=0.143, p=n.s; Fig 2B).Effects of nuclear volume were taken into account. These findings indicate that maternalising the paternal domain and paternalising the maternal domain, do not result in the expected shift in localisation. The discordance between of the findings in Figure 2A and Figure 2B experiments suggests that the epigenotype switch in the two imprinting models is not reflecting the parental origin of wildtype chromosomes in terms of subnuclear location. However, these data do not consider the transcription of imprinted loci in these models.

### Stochastic or developmental repression of the maternally expressed *Gtl2/Meg3* gene correlates with a shift of alleles towards the nuclear periphery in ES cells

While *Dlk1* is not expressed in ES cells, *Gtl2/Meg3* is strongly expressed. Though all maternally inherited alleles of the *Gtl2/Meg3* gene have the potential to express the gene, it is not active in all cells. It is not known whether this effect is stochastic or regulated. Gene expression can be fine-tuned by transcription on-off cycles, sometimes called “bursting”, which could reflect genes moving in and out of transcription factories (3). Furthermore, even in the pluripotent state, some ES cells will start to differentiate, with concurrent changes in gene expression, and eventually will stop dividing in ES culture conditions. Since we cannot distinguish these possibilities in our experiments, we refer to *Gtl2/Meg3*-non-expressing maternal alleles as “stochastic or developmental repression”. Stochastic or developmental repression of the canonically active maternal allele of *Gtl2/Meg3* was observed in 29.5% of WT ES cells and 32.3% of patRepKO ES cells (Supplementary Fig. S4). We took advantage of this property to test whether active and repressed *Gtl2/Meg3* alleles differed in their intranuclear localisation using RNA FISH. We used a logistic regression approach to test if distance to the periphery predicted the expression state of *Gtl2/Meg3* (after controlling for nuclear volume variation). In three biological replicates each of patrepKO ES cell lines in which the single maternally inherited *Meg3* allele has the potential to be either expressed or not, we found no difference in localisation regardless of the expression of *Gtl2/Meg3* (z=1.059, p=n.s; Fig. 3A).

**Figure 3:**
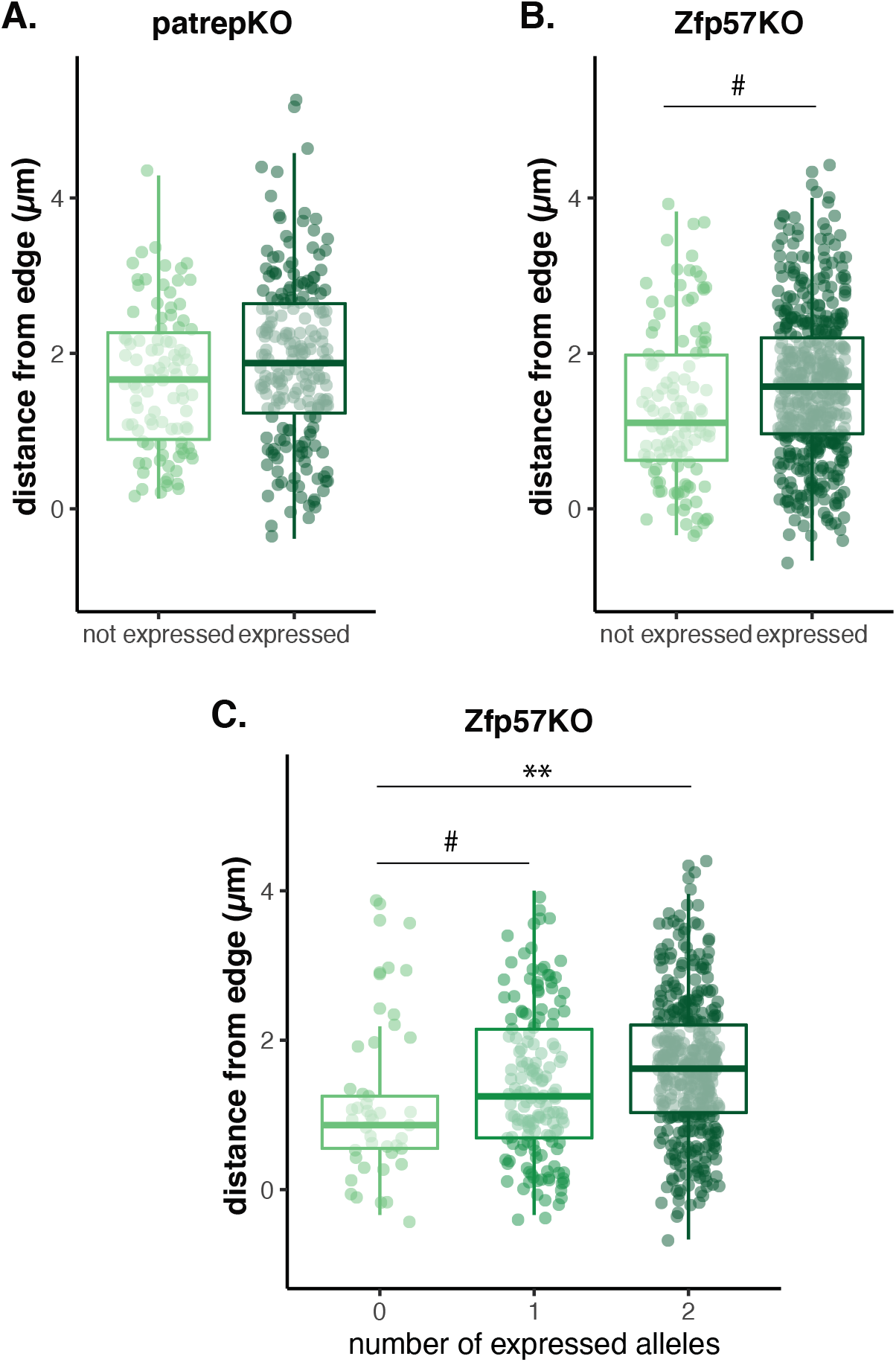
Distribution of alleles depending on *Gtl2/Meg3* gene expression.

(**A**) There was no relationship between localisation and *Gtl2/Meg3* expression from patrepKO ESCs. (**B**) Maternalized alleles from Zfp57KO ES cells were marginally closer to the edge when *Gtl2/Meg3* was non-expressed (#*p*<0.1), (**C**) Maternal(ized) alleles split by number of alleles that were expressed. Alleles that were expressed alone in the cell were marginally farther from the periphery than alleles that were not expressed. However, alleles that had both alleles expressed within the cell were significantly further away from the periphery (*p*<0.1, \**p*<0.05).

However, since Zfp57KO ES cells have twice as many maternal(ized) alleles as patrepKO ES cells, we reasoned that experiments with these cell lines would have greater power to examine the relationship between distance and expression. Using the same statistical methodology and same numbers of replicates, we found a non-significant trend (estimate =0.927, p=0.0721, Fig. 3B) with non-expressed alleles being marginally closer to the periphery compared to expressed alleles. As before, nuclear volume was taken into account. Together these findings indicate insignificant effects of repression on localisation relative to the nuclear periphery. We therefore focused on expressed alleles to consider whether the expression rather than repression might predict localisation.

### Gene expression predicts localisation better than parental epigenotype

Using Zfp57KO, we can measure whether biallelic expression could predict distance from the nuclear border better than monoallelic expression. Using a multiple linear mixed model approach we find that there is a marginal increase in distance when only one allele within the cell is expressed (t=1.837, p=0.067, estimate (slope) = 0.175) but alleles move significantly further from the nuclear border when both alleles are expressed (t=1.989, p<0.05, estimate (slope) =0.185; Fig 3C). This suggests that it is expression rather than parental epigenotype or repression that confers subnuclear localisation. Consistent with this and as shown in Fig 4A, expressed alleles are more likely to be greater than 0.5μm away from the periphery (*χ*^2^=3.0713, df=1, p=0.08). To examine this at higher resolution, we plotted the regional location after dividing the nucleus into thirds by volume (outer, middle, inner) as has been described previously (19). Results suggest that repressed alleles are not preferentially located to the periphery, rather, active alleles are enriched away from the periphery (*χ*^2^=14.206, df=2, p<0.001; Fig 4B).

**Figure 4:**
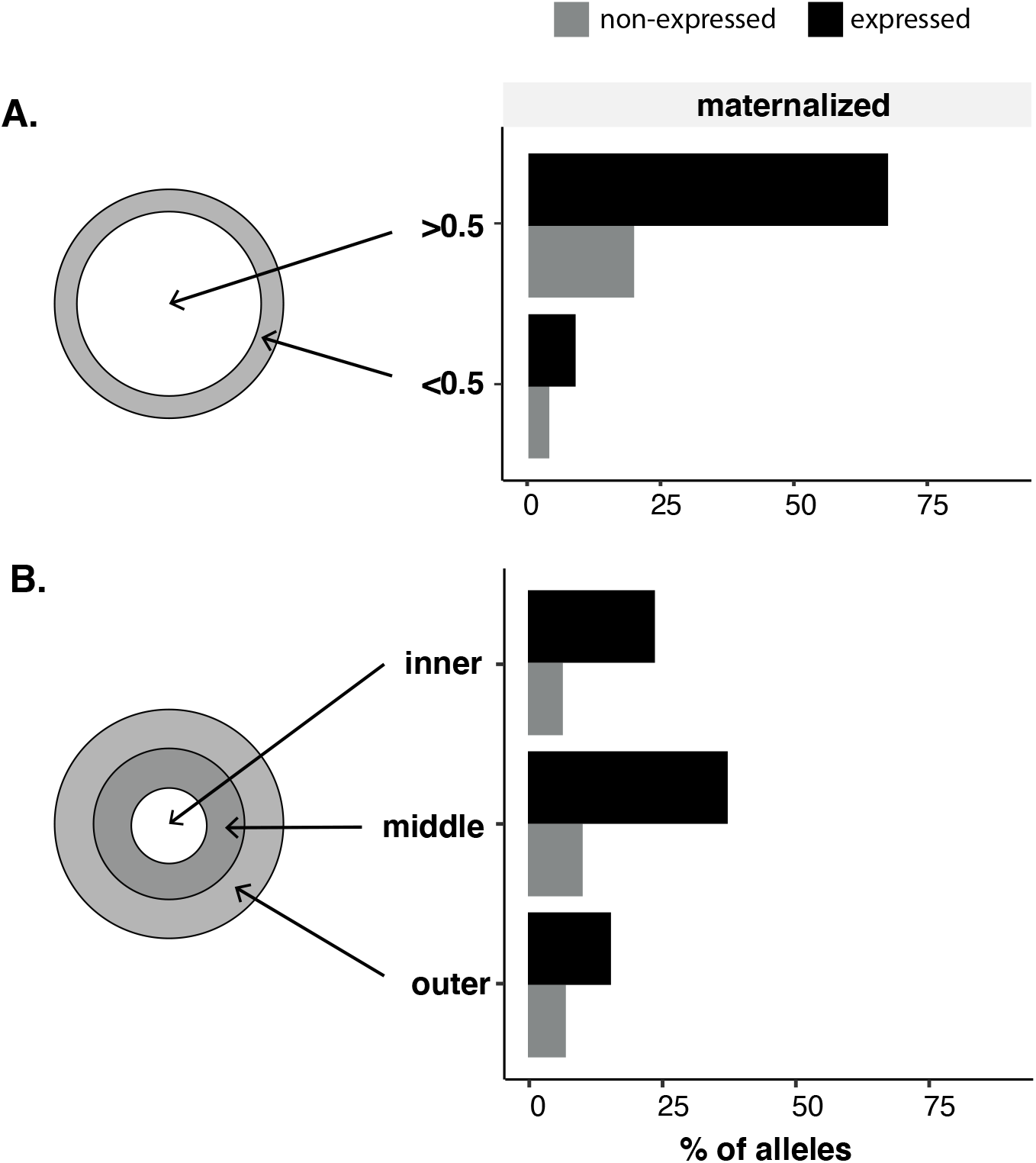
*Gtl2/Meg3*-non-expressing maternal(ized) alleles are not enriched at the nuclear periphery.

(**A**) *Gtl2/Meg3* expressing chromosomes (patrepKO and Zfp57KO) alleles show a significant shift away from the nuclear periphery compared to non-expressed alleles. (**B**) If the allele distribution is depicted as relative to the nuclear radius, the shift is still visiblewith a greater proportion of *Gtl2/Meg3* expressed alleles in inner and middle areas.

## Discussion

We performed 3D RNA-DNA FISH measurements in a total of 18 mouse ES cell lines and observed a very small but overall highly significant effect for *Gtl2/Meg3*-expressing alleles of the *Dlk1-Dio3* imprinted region to be localized towards the nuclear interior and for non-expressing alleles to be localized towards the nuclear periphery. As has been shown before (19), paternal alleles, which do not express *Gtl2/Meg3*, localize overall closer to the periphery than maternal alleles, from which *Gtl2/Meg3* is expressed in 70.5% of WT ES cells. Differences in localisation between paternalized and maternalized allelles (i.e., IGDMRKO *vs* ZFP57KO) are only evident when expression state is considered. In other words, expressed *Gtl2/Meg3* alleles are positioned further away from the periphery than non-expressed alleles. Furthermore, the position within the nucleus is directly related to the number of alleles being expressed within a given cell, as evident through the analysis of cells expressing zero, one or two alleles.

The effect size of the localisation difference between expressing and non-expressing alleles in our study is much smaller than that previously described (19). Three notable differences between our work and the previous study might explain this: 1) Kota et al. used *Gtl2/Meg3* expression as a read-out for parental origin while we included an independent DNA marker and genetic approaches allowing us to distinguish and quantify expressing and non-expressing maternal chromosomes. 2) Kota et al. used one biological replicate hence would not have been able to take into account the variability that is evident between lines. 3) Our edge measurements utilise a DNA marker located 400kb away from the interrogated gene which might result in lower data resolution (supplementary Fig. S1). Both genetic locations lie, however, within the same TAD in ES cells (23,24) and show frequent overlap in a double probe DNA FISH experiment using probes against the L1 repeat and the *Gtl2/Meg3* gene (Supplementary Fig. S1 and S5). Our findings reflect the importance of independently distinguishing parental chromosomes and expression and then determining the contribution that each makes to subnuclear localisation.

Mouse ES cells show an unusual cell cycle distribution where a majority of cells (75%) will be in S-phase in a growing cell population (28). S-phase can potentially have an influence on gene expression and cell cycle heterogeneity could thus be a source of variability between biological replicates. However, when comparing nuclear sizes (as a proxy for cell-cycle differences) of expressing and non-expressing alleles as well as biological replicates, some small variation in nuclear size was evident in some instances (Supplementary Fig. S6). We therefore included nuclear volume as a covariate in all of our analyses to correct for any minor fluctuations in nuclear volume. Hence, all of our estimates of the differences in allelic distribution control for minor changes in nuclear volume.

Though data from all our ES cell lines show a significant expression-dependent effect on subnuclear localisation, the effect size is extremely small with *Gtl2/Meg3*-non-expressing alleles being on average only 170 nm closer to the nuclear periphery than *Gtl2/Meg3*-expressing alleles. Indeed, most expressed alleles can be more broadly categorised as being away from the periphery (or outer most region) though not extremely central to the nucleus. It is interesting to consider whether such a small effect is biologically meaningful. Since we did not observe a significant enrichment of inactive alleles in the immediate vicinity of the nuclear envelope, inactivating interactions with the nuclear lamina are unlikely to play a major role in the observed allele distribution. Notably, *Gtl2/Meg3* expression was regularly found right at the edge of DAPI staining and occasionally outside of it, probably representing areas of low density chromatin at the very periphery of nuclei (see methods section for details). Although it can be envisioned that in some cases *Gtl2/Meg3* expression might have been inhibited by transient nuclear envelope interactions and had already moved away from the periphery at the time of the experiment, such alleles should still show up as peripheral enrichment since mobility of individual loci during interphase is usually limited to about 1 μm and it has been shown that large scale changes in localization require cell division (1,29). A more likely explanation is that the observed pattern of allele localisation is due to some activating effect of structures at the nuclear interior. This is consistent with the increase in internalisation observed when two alleles are expressed compared to one in a given cell. Transcription is known to happen preferentially at sites where the transcriptional machinery and active genes are clustered into so-called transcription factories (3,4). Such factories are found in open chromatin at the borders of chromosomal territories and it seems logical that they would be biased to be more frequent at the more transcriptionally favourable nuclear interior. Therefore, one possible explanation for the observed non-enrichment at the nuclear periphery and the small effect size of our expression-localisation correlation could be that *Gtl2/Meg3*-non-expressing alleles are evenly distributed within the space they can take up around the chromosome territory, independent of nuclear envelope interactions, while *Gtl2/Meg3*-expressing alleles need to be localized at transcription factories which are biased to be at the interior. It would be interesting to test this idea by co-localising expressing and non-expressing alleles with components of transcriptional factories. Nevertheless, our use of multiple biological replicates in multiple genetic models highlights the wide range of positions that an allele can take within the nucleus regardless of expression status.

For all correlations of nuclear localisation and expression, the question of cause and consequence arises. Is peripheral localisation used as a means to fine-tune gene expression by making the co-localisation with transcription factories more or less likely? The absence of enrichment of inactive alleles at the nuclear periphery does not argue for such a mechanism. Kota and colleagues have shown that position changes of the *Dlk1-Dio3* region are local and do not involve gross changes in chromosome territory (19). Similarly, our Zfp57 mutants show that local and purely epigenetic changes are sufficient to cause a shift in localisation at the same time as inducing *Gtl2/Meg3* gene expression. This argues that a nuclear envelope anchoring mechanism acting at a distance is unlikely to be involved. A local anchoring mechanism on the other hand should show up as peripheral enrichment which we do not observe. We therefore suggest that the small shift in peripheral localisation that we observe is a consequence, rather than a cause of expression changes and that the more functionally relevant aspect is the interior location of expressed alleles which is influenced by the number of alleles being expressed.

## Methods

### Cell culture

Mutant and corresponding control ES cells were generated from single blastocyst using feeder-free-based 2i LIF culture conditions (N2B27, Stem Cell Sciences) (30). In brief, morula embryos were collected from pregnant female mice and cultured in KSOM with 2i inhibitors, and then each blastocyst was genotyped using trophectoderm after immunosurgery. *del*^L1rep^ mice have an *albino* C57BL/6J background (BL6) (20) and were crossed to *albino* BL6 mice to obtain WT, maternal and paternal heterozygotes. IG-DMR heterozygous females (27) were mated with BL6 males to obtain IG-DMR maternal KO and control WT morula embryos. *Zfp57* zygotic KO ES cells and corresponding control ES cells were generated in a previous study (26). The research was conducted in accordance with UK Home Office Animals Scientific Procedures Act, project licence 80/2567. We observed unexpectedly low expression levels of *Gtl2/Meg3* in two of five *Zfp57KOES* cell lines and therefore excluded these two cell lines from these studies. For our analysis of ES cells of all genotypes, we used data from the three biological replicates that showed the expected and similar proportions of non-expressing, monoallelically expressing, and biallelically expressing cells (Supplementary Fig. S2).

### Fluorescence in situ hybridisation

We generated fluorescent probes for detection of the LINE repeat (BAC bMQ-177C10, obtained from CHORI) and nascent *Gtl2/Meg3* transcripts (fosmid WIBR1-2686H19, a kind gift from the Heard lab, Paris) by first amplifying plasmid mini preps with the Illustra TempliPhi Large Construct Kit (GE Healthcare) and subsequently labelling 2 μg of the amplification product by nick translation using Green UTP or Red UTP (Abbott Molecular) according to the manufacturer’s instructions. Sequential RNA and DNA 3D fluorescence in situ hybridisation (FISH) was performed as previously described (31). However, ES cells were grown on laminin-coated coverslips prior to fixing to ensure well-spread monolayer growth and nuclei were counterstained with 0.2 μg/ml DAPI after hybridisation and washes. 3D image stacks of 10-15 positions per coverslip were acquired using a Carl Zeiss Axiovert 200M microscope with a 63x/1.25 Plan Apochromat objective taking 100 image planes per image stack at a z spacing of 0.15 μm. For double probe DNA FISH the same protocol was used as in sequential FISH and the Gtl2/Meg3 fosmid probe was used to detect the gene rather than nascent RNA.

### Image analysis

DNA FISH image stacks were post-processed using Huygens Professional deconvolution software (version 14.10.1p8, SVI, The Netherlands) and 3D measurements were taken using Fiji (32). We developed a Fiji script that enabled us to generate binary representations of DAPI-stained nuclei and FISH signals and to take automated 3D measurements of and between these objects calling Fiji’s 3D manager function (Fig. 1C). If FISH signals were duplicated due to DNA replication, we used the mean of both measurements for the analysis. Due to the nature of the binarization process using DAPI staining of chromatin, binary images of nuclei sometimes differed slightly in shape from what was seen by eye in the original image because, depending on image background levels or closeness of neighbouring nuclei, non-dense chromatin regions were interpreted as background by the algorithm. This led to a minority of DNA FISH signals (2.8%) being outside the nuclear volume. To avoid bias against peripherally located signals, we included such signals as negative distance measurements. In very rare cases the algorithm generated a bay-like background area around a FISH signal in a non-dense chromatin area, in a way that our standard automated measurements would lead to false results. Again, to avoid biasing against peripheral signals as well as inaccurate manual measuring, in these rare cases (0.3%) we used an alternative algorithm that approximates the nuclear volume by fitting an ellipse around the original shape and used this shape for distance measurements. For relative distance measurements, we used the “radiusCen” measurement of Fiji’s 3D manager, which measured the distance between the centre of the binary representation of the nucleus and its border through the centre of the binary representation of the FISH signal, as local nuclear radius. Relative distances were calculated as absolute distance / local nuclear radius. In parallel to DNA FISH analysis, the expression state of the *Gtl2/Meg3* gene was evaluated from original (non-deconvolved) nascent RNA FISH image stacks for each allele as either expressed (FISH signal above background level) or non-expressed (no signal).

### Statistical methodology

All analyses used a linear regression model framework using nuclear size as a covariate (to control for any volume effects on expression and/or localisation) using base R regression functions (33). Given that some cells had measurements for multiple alleles, where appropriate, we used mixed effects linear regression models using the lme4 and lmerTest packages in R (34,35) to allow for random intercepts (of unique cells) which effectively controls variability arising from the repeated measurement of unique cells (36). Data handling and visualisation in R was carried out using the ‘tidyverse’ packages (37). Chi-squared tests for differences in nuclear volume distributions were calculated using base chi.sq functions in R.

## Acknowledgements

We would like to thank Richard Butler at the Gurdon Institute Imaging Facility, Cambridge, for help with writing Fiji macros for 3D image analysis, Edith Heard, Simao daRocha and Luca Giorgetti at the Institute Pierre et Marie Curie, Paris, for helpful advice with FISH methods, and David Glover and George Tsolovsky at the Department of Genetics, Cambridge, for microscopy support. This work was funded by a DFG research fellowship NE 1959/1-1 to LCH and grants from the Wellcome Trust and MRC to ACFS. The authors declare no conflict of interest.

## Authors contributions

LCH and ACFS conceived and designed the experiments. LCH and FLD conducted the FISH experiments and LCH and RM conducted the data analysis. NT generated and tested the ES cell lines. LCH, RM and ACFS wrote the paper.Figure legends

## Supplementary Information

**Supplementary Figure S1:** UCSC (https://genome.ucsc.edu/)-screenshot of the Dlk1-Dio3 imprinted region in mouse mm9 showing the exact positions of RNA (Meg3) and DNA (L1 repeat) FISH probes, as well as the genomic distances between them (inner, central and outer distance). The exact position of the LINE1 repeat deletion (20) in relation to the repeat probe and the UCSC RepeatMasker are also shown. The top panel shows ES cell TADs from Schoenfelder and colleagues (Bab_ESC) and Dixon and colleagues (Ren_ESC) loaded as custom tracks (23,24)

**Supplementary Figure S2:** Plots of the distance from the nuclear border relative to *Gtl2/Meg3* expression in individual biological replicates of all KO and WT groups measured in the study. We find no significant effect of replicate on overall distance measures.

**Supplementary Figure S3:** Additional examples of gene expression in the five ES cells genotypes at larger magnifications (compare figure 1). *Gtl2/Meg3* probe stained images are from nascent RNA FISH and L1 Repeat probe stained images are from subsequent DNA FISH. The FISH images represent maximum projections of 10-30 central z planes of acquired image stacks. Blow-ups of framed regions are shown on the right. Scale bars represent 50 μm.

**Supplementary Figure S4:** Numbers of cells per cell line with no, monoallelic, or biallelic *Gtl2/Meg3* expression.

**Supplementary Figure S5:** Double probe DNA FISH in WT ES cells (ES cell line WT8). The images show only one z image plane. All three locations, at which the L1 repeat signal and the *Gtl2/Meg2* gene signal have their centres roughly in the same z image plane, are shown as blow-ups on the right. The distance between the signal centres is between 0.25 and 0.36 μm in these three examples.

**Supplementary Figure S6**: The density of nuclear sizes does not vary considerably between *Gtl2/Meg3*-expressing and -non-expressing ES cells and therefore does not hint towards a specific down- or upregulation of gene activity during s-phase.

**Supplementary Table 1**: CSV file of raw data and metadata from all experimental cell lines.

## Notes

### Competing Interest Statement

The authors have declared no competing interest.

## References

1. van Steensel B, Belmont AS. Lamina-Associated Domains: Links with Chromosome Architecture, Heterochromatin, and Gene Repression. Vol. 169, Cell. 2017.

2. Cremer T, Cremer C. Chromosome territories, nuclear architecture and gene regulation in mammalian cells. Vol. 2, Nature Reviews Genetics. 2001. p. 292–301.

3. Osborne CS, Chakalova L, Brown KE, Carter D, Horton A, Debrand E, et al. Active genes dynamically colocalize to shared sites of ongoing transcription. Nature Genetics. 2004;36(10):1065–71.

4. Nguyen HQ, Bosco G. Gene Positioning Effects on Expression in Eukaryotes. Annual Review of Genetics. 2015;49(1).

5. Shachar S, Misteli T. Causes and consequences of nuclear gene positioning. Journal of Cell Science. 2017;130(9).

6. Guelen L, Pagie L, Brasset E, Meuleman W, Faza MB, Talhout W, et al. Domain organization of human chromosomes revealed by mapping of nuclear lamina interactions. Nature. 2008;453(7197):948–51.

7. Kind J, Pagie L, Ortabozkoyun H, Boyle S, De Vries SS, Janssen H, et al. Single-cell dynamics of genome-nuclear lamina interactions. Cell. 2013;153(1):178–92.

8. Casolari JM, Brown CR, Komili S, West J, Hieronymus H, Silver PA. Genome-wide localization of the nuclear transport machinery couples transcriptional status and nuclear organization. Cell. 2004;117(4):427–39.

9. Kalverda B, Pickersgill H, Shloma V V., Fornerod M. Nucleoporins Directly Stimulate Expression of Developmental and Cell-Cycle Genes Inside the Nucleoplasm. Cell. 2010;140(3):360–71.

10. Capelson M, Liang Y, Schulte R, Mair W, Wagner U, Hetzer MW. Chromatin-Bound Nuclear Pore Components Regulate Gene Expression in Higher Eukaryotes. Cell. 2010;140(3):372–83.

11. Vaquerizas JM, Suyama R, Kind J, Miura K, Luscombe NM, Akhtar A. Nuclear pore proteins Nup153 and megator define transcriptionally active regions in the Drosophila genome. PLoS Genetics. 2010;6(2).

12. Finlan LE, Sproul D, Thomson I, Boyle S, Kerr E, Perry P, et al. Recruitment to the nuclear periphery can alter expression of genes in human cells. PLoS Genetics. 2008;4(3).

13. Reddy KL, Singh H. Using molecular tethering to analyze the role of nuclear compartmentalization in the regulation of mammalian gene activity. Methods. 2008;45(3):242–51.

14. Zullo JM, Demarco IA, Piqué-Regi R, Gaffney DJ, Epstein CB, Spooner CJ, et al. DNA sequence-dependent compartmentalization and silencing of chromatin at the nuclear lamina. Cell. 2012;149(7):1474–87.

15. Dialynas G, Speese S, Budnik V, Geyer PK, Wallrath LL. The role of Drosophila Lamin C in muscle function and gene expression. Development. 2010;137(18):3067–77.

16. Kumaran RI, Spector DL. A genetic locus targeted to the nuclear periphery in living cells maintains its transcriptional competence. Journal of Cell Biology. 2008;180(1):51–65.

17. Ferguson-Smith AC. Genomic imprinting: the emergence of an epigenetic paradigm. Nature Reviews Genetics. 2011;12(8):565–75.

18. McEwen KR, Ferguson-Smith AC. Distinguishing epigenetic marks of developmental and imprinting regulation. Epigenetics and Chromatin. 2010;3(1).

19. Kota SK, Llères D, Bouschet T, Hirasawa R, Marchand A, Begon-Pescia C, et al. ICR noncoding RNA expression controls imprinting and DNA replication at the Dlk1-Dio3 domain. Developmental Cell. 2014;31(1):19–33.

20. Soares ML, Edwards CA, Dearden FL, Ferron SR, Curran S, Corish J, et al. Targeted deletion of a 170 kb cluster of LINE1 repeats: implications for regional control. Genome research. 2018;gr.221366.117.

21. Peric-Hupkes D, Meuleman W, Pagie L, Bruggeman SWM, Solovei I, Brugman W, et al. Molecular Maps of the Reorganization of Genome-Nuclear Lamina Interactions during Differentiation. Molecular Cell. 2010;38(4):603–13.

22. Sun B, Ito M, Mendjan S, Ito Y, Brons IGM, Murrell A, et al. Status of genomic imprinting in epigenetically distinct pluripotent stem cells. Stem Cells. 2012;30(2):161–8.

23. Schoenfelder S, Furlan-Magaril M, Mifsud B, Tavares-Cadete F, Sugar R, Javierre BM, et al. The pluripotent regulatory circuitry connecting promoters to their long-range interacting elements. Genome Research. 2015;

24. Dixon JR, Selvaraj S, Yue F, Kim A, Li Y, Shen Y, et al. Topological domains in mammalian genomes identified by analysis of chromatin interactions. Nature. 2012;485(7398):376–80.

25. Li X, Ito M, Zhou F, Youngson N, Zuo X, Leder P, et al. A Maternal-Zygotic Effect Gene, Zfp57, Maintains Both Maternal and Paternal Imprints. Developmental Cell. 2008;15(4):547–57.

26. Shi H, Strogantsev R, Takahashi N, Kazachenka A, Lorincz MC, Hemberger M, et al. ZFP57 regulation of transposable elements and gene expression within and beyond imprinted domains. Epigenetics Chromatin. 2019 Aug 9;12(1):49.

27. Lin SP, Youngson N, Takada S, Seitz H, Reik W, Paulsen M, et al. Asymmetric regulation of imprinting on the maternal and paternal chromosomes at the Dlk1-Gtl2 imprinted cluster on mouse chromosome 12. Nature Genetics. 2003;35(1):97–102.

28. Savatier P, Lapillonne H, Jirmanova L, Vitelli L, Samarut J. Analysis of the Cell Cycle in Mouse Embryonic Stem Cells. Embryonic Stem Cells. 185(3):27–33.

29. Kind J, Pagie L, De Vries SS, Nahidiazar L, Dey SS, Bienko M, et al. Genome-wide Maps of Nuclear Lamina Interactions in Single Human Cells. Cell. 2015;163(1).

30. Ying Q-L, Ying Q-L, Wray J, Wray J, Nichols J, Nichols J, et al. The ground state of embryonic stem cell self-renewal. Nature. 2008;453(May):519–23.

31. Giorgetti L, Galupa R, Nora EP, Piolot T, Lam F, Dekker J, et al. Predictive polymer modeling reveals coupled fluctuations in chromosome conformation and transcription. Cell. 2014;157(4):950–63.

32. Schindelin J, Arganda-Carreras I, Frise E, Kaynig V, Longair M, Pietzsch T, et al. Fiji: an open-source platform for biological-image analysis. Nature Methods. 2012;9(7):676–82.

33. R Core Team. R: A Language and Environment for Statistical Computing [Internet]. Vienna, Austria: R Foundation for Statistical Computing; 2019. Available from: https://www.R-project.org/

34. Bates D, Mächler M, Bolker B, Walker S. Fitting Linear Mixed-Effects Models Using lme4. J Stat Soft [Internet]. 2015 [cited 2020 Jan 3];67(1). Available from: http://www.jstatsoft.org/v67/i01/

35. Kuznetsova A, Brockhoff PB, Christensen RHB. lmerTest Package: Tests in Linear Mixed Effects Models. J Stat Soft [Internet]. 2017 [cited 2020 Jan 3];82(13). Available from: http://www.jstatsoft.org/v82/i13/

36. Gelman A, Hill J. Data Analysis Using Regression and Multilevel/Hierarchical Models [Internet]. Cambridge: Cambridge University Press; 2006 [cited 2020 Jan 3]. Available from: http://ebooks.cambridge.org/ref/id/CBO9780511790942

37. Wickham H, Averick M, Bryan J, Chang W, McGowan L, François R, et al. Welcome to the Tidyverse. JOSS. 2019 Nov 21;4(43):1686.

